# MiRNA Locker: A Modularized DNA Assembly As miRNA Inhibitors

**DOI:** 10.1101/2025.01.10.632138

**Authors:** Lingyun Zhu, Lu Min, Chu-shu Zhu, Xiaomin Wu, Wenying Li, Jiaxin Ma, Yilin Lei, Changsong Gao, Xinyuan Qiu, Chuanyang Liu

## Abstract

MicroRNAs (miRNAs) are vital post-transcriptional regulators that govern key cellular processes such as proliferation, migration, and apoptosis. Current loss-of-function approaches, including chemically modified antisense oligonucleotides (ASOs), face significant challenges, including high costs, limited scalability, and off-target effects. To overcome these limitations, we developed “miRNA Locker”, a novel miRNA inhibition platform created using the Overlapped Oligo Assembly (OOA) method. This innovative platform constructs highly stable dumbbell-shaped single-stranded DNA structures, offering improved target specificity, scalability, and cost-effectiveness. Using miR-214 as a proof-of-concept target, we demonstrated that miRNA Lockers effectively bind Argonaute-miRNA complexes, reduce miRNA levels, and induce downstream changes in gene expression and cellular phenotypes, surpassing the performance of commercial antagomirs. Furthermore, applying miRNA Lockers to miR-654 validated the regulatory role of miR-654 in modulating RNF8 expression and promoting epithelial-mesenchymal transition (EMT) in lung cancer cells. Our results highlight the potential of miRNA Lockers as a versatile tool for studying miRNA function and advancing miRNA-based therapies.

**Graphical Abstract:** 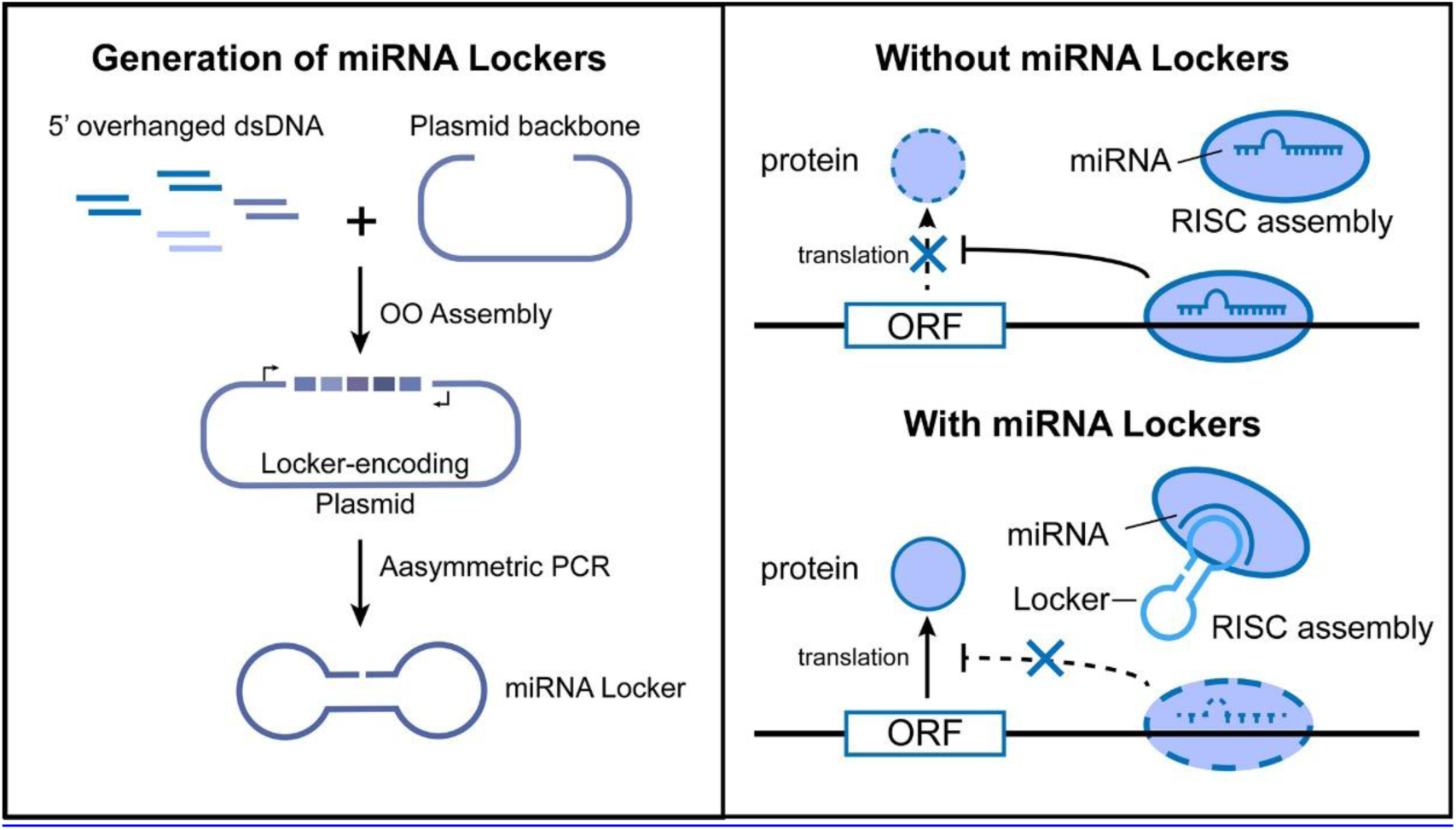

## Introduction

MicroRNAs (miRNAs) are small non-coding RNAs 18-24 nucleotides in length that have been proven to play important roles on post-transcriptional regulation of the gene expression^1^. Up to now, over 2000 miRNAs have been identified or predicted from human tissues or cells^2^. Mature miRNAs are mainly existing in cytoplasm and, together with Argonaute (Ago) family of protein, packed into a protein complex known as RNA-induced silencing complex (RISC)^3^. Base-paring to 3’ untranslated region (3’ UTR) of target mRNA, miRNA would induce translational repression or mRNA degradation due to the endonucleolytic activity of Ago, thus regulating gene expression and participants in many pivotal biological processes, including cell proliferation, differentiation, migration and apoptosis^4–6^. The 2024 Nobel Prize in Physiology or Medicine highlighted the fundamental discoveries of miRNA-mediated gene regulation, emphasizing their critical roles in cellular homeostasis and therapeutic potential. To understand the regulatory mechanism of miRNA, miRNA gain of function and loss of function tools are necessary. Thus, methods for inhibiting functional microRNAs *in vitro* and *in vivo* is wildly needed in miRNA researches^7–9^.

Presently, loss-of-function phenotypes are mainly induced by means of chemically modified antisense oligonucleotides(ASO). Chemically modified antisense oligonucleotides includes 2’O-methyl, 2’O-methoxyethyl, locked nucleic acid (LNA) and others, which aims to pair with and block mature microRNAs through strictly sequence complementarity^10^. Antagomirs are a group of chemically modified antisense oligonucleotides, which are readily available tools wildly used for endogenous miRNA inhibition^3^. To improve the stability and miRNA binding affinity, different modifications are introduced to antagomir and optimized to achieve high fidelity, low toxicity, and improved stability. Although there are many advantages of ASO, limited scalability and off-target effects still limit the extensive usage of ASO in miRNA loss of function study.

Apart from ASO, two other approaches have been reported to be utilized in miRNA loss of function researches. One is called “miRNA sponge”, which stands for a class of competitive inhibitor of miRNA carrying multiple binding sites of miRNA^11^. Another is termed “miRNA mask” ^12^, which uses oligonucleotides to be perfectly complementary to mRNA, like a mask covering miRNA binding sites. Therefore, miRNA cannot bind to mRNA and subsequently induce miRNA-mRNA interaction. However, miRNA inhibition using miRNA sponge or miRNA mask are not widely used for miRNA loss of function researches^10^. Notably, previous reports have demonstrated that miRNA can efficiently bind to single-strand DNA in a base-matching manner. Deng et al. also showed that non-modified DNA with dumb-bell-shaped secondary structure and miRNA binding sites on the loop region presents high miRNA binding affinity and improved stability^13^. Hence, we anticipate that dumb-bell-shaped DNA may serve as miRNA inhibitors as well.

In this paper, we demonstrated a dumb-bell-shaped DNA-based design of miRNA inhibitor to achieve low-cost and high scalability with high inhibitory effect simultaneously. Our miRNA inhibitor, named miRNA locker, can be easily assembled using modularized DNA parts from a set of chemically synthetic oligo DNA libraries. This study aims to evaluate their inhibitory effectiveness compared to existing miRNA inhibitors and explore their potential applications in miRNA research and therapeutic interventions.

## Results

### Modular Design and assembly of miRNA Lockers

MiRNA Lockers are unsealed dumbbell shaped single-stranded DNAs (ssDNA) that is capable of binding specific miRNAs **(Figure 1a)**. The double-stranded region, named as the supporting module, serves as a “handle” to improve the stability of Locker. The loop regions, known as the functional modules, are designed to bind specific miRNAs through Watson-Crick base pairing. Intervals are inserted between modules for the convenience of the assembly procedure. Though short miRNA Lockers can be directly synthesized, it is noticeable that the cost of direct synthesis raise rapidly as the length of the Locker increases. Hence, a low-cost and rapid assemble protocol is needed. Due to the difficulties in ligating ssDNAs, our strategy is to assemble double-strand DNA (dsDNA) containing the Locker strand and its complementary strand for the first step, then use techniques such as asymmetric PCR to generate ssDNAs^14^.

**Figure 1.**
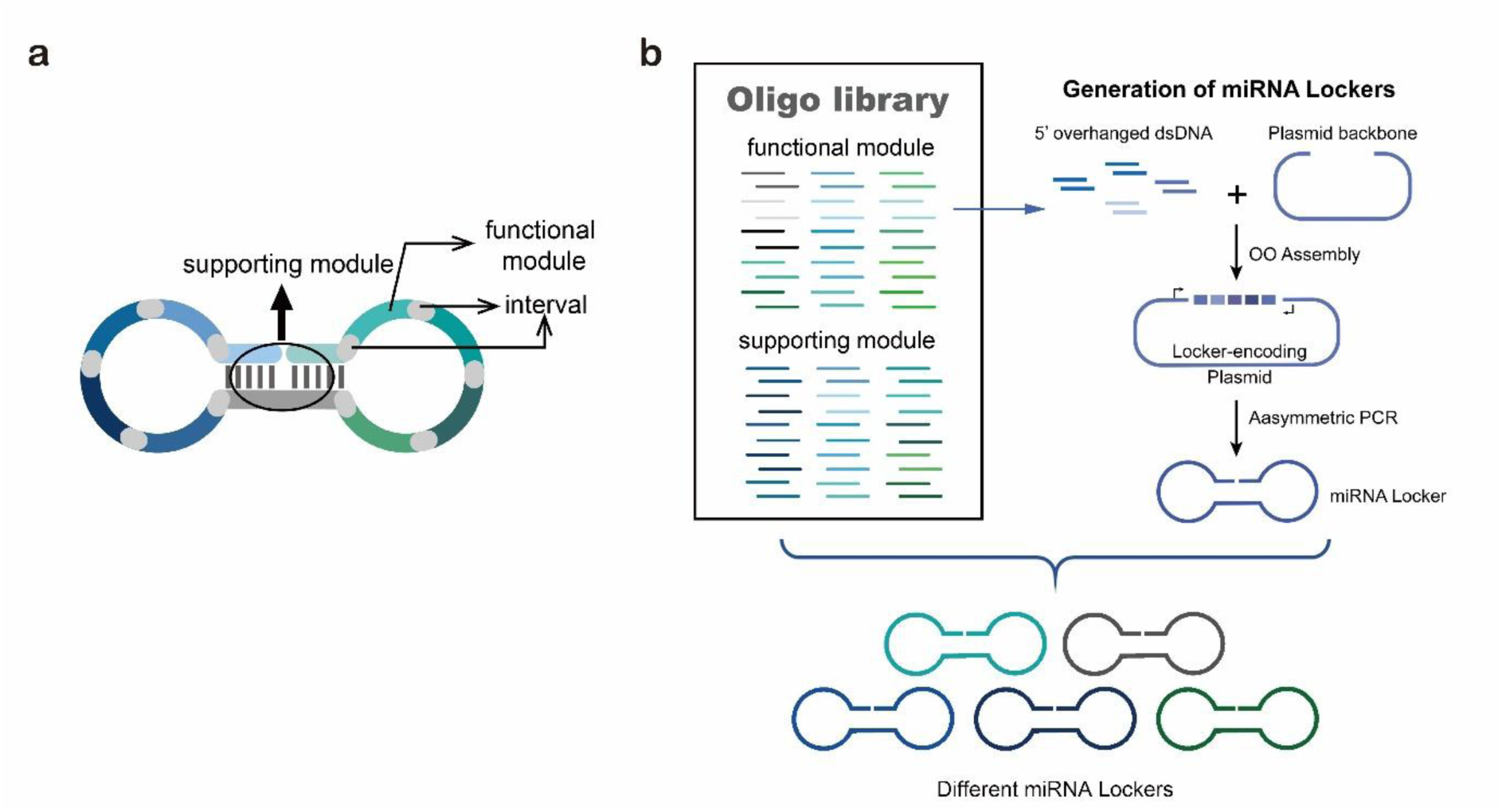
Modular design of miRNA Lockers via OOA assembly. (a) Design and structural diagram of miRNA locker. (b) The construction and assembly of miRNA lockers.

Noticing that the Locker-containing dsDNA can be divided into several modularized DNA parts according to the structure of the Locker, we modified a previously reported oligo-linker mediated assembly (OLMA) method to assemble the Locker-containing dsDNA ^15^. In our new method, named as **Overlapped Oligo Assembly (OOA) method**, modularized DNA parts is designed into the chemically synthesized DNA oligo, and overlapped 5’-overhangs are added for assembly. After that, dsDNAs can be obtained by annealing of two complementary single-strand oligonucleotides, then phosphorylated by T4 Polynucleotide Kinase to facilitate the following ligation reaction. T4 DNA ligase is use for their ligation onto pSB1C3 plasmid backbone. High-throughput one-step assemble of multiple short dsDNA nucleotides can therefore be achieved, and the correct assembling order is guaranteed by using identical overlap sequences among oligos, serving as zipcodes. The zipcodes are introduced as the intervals in a miRNA Locker **(Figure 1b and Figure 2a)**. As a primary validation, four experimental cases were constructed to examine the scalability, reliability and efficiency of the OOA method. Specifically, Locker A was designed with two identical functional modules to target miR-214, Locker B contained two different functional modules to target miR-214 and miR-654 at the same time, Locker C contained three identical modules to target miR-214, and Locker D contained four different modules targeting let-7a, miR-16, miR-195 and miR-184 **(Figure 2b, left panel)**. Nucleotides chosen from the library encoding the supporting modules and functional modules of each of the four Locker designs were assembled using OAA method and transformed into *E.coli* DH5α chassis. Assemblies were validated via sequencing using universal primers for pSB1C3 backbone. Results showed >50% success rates in each case, with few mutations identified. **(Figure 2b, middle panel)**.

**Figure 2.**
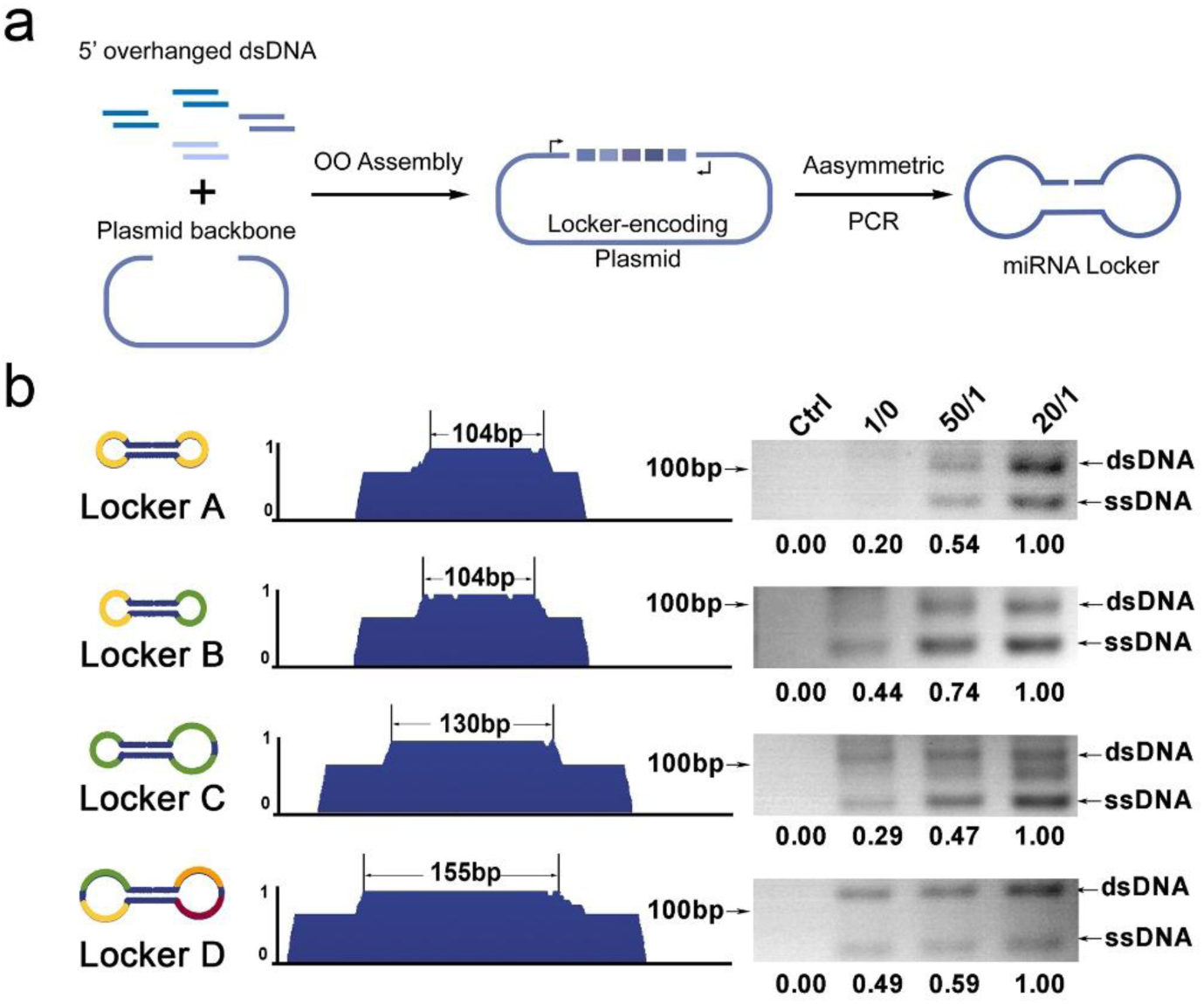
Generation of miRNA Lockers with OAA assembly and asymmetric PCR. (a) Schematic representation of the Locker generating procedure. (b) Left panel: Design of four testing cases for the validation of the locker-generating procedure. Different miRNA binding modules were shown by different colors; middle panel: Sequencing validation for the ds DNA assemblies encoding four miRNA lockers; right panel: electrophoresis showing the production of four miRNA lockers by asymmetric PCR.

Afterwards, to generate single strand miRNA Lockers, miRNA-Locker-coding region was digested from the plasmid and then used as the template for asymmetric PCR. To optimize the PCR protocol, primers in ratios of 20/1, 50/1 and 1/0 were tested. The asymmetric PCR products were analyzed through 3% agarose gel electrophoresis. As showed in the right panel of **Figure 2b**, bands showing ssDNA products were observed in all four cases when 20:1, 50:1 and 1:0 primer ratios were applied. Gray-scale analysis normalized with 20:1 group showed that the ssDNA amplification signal in 20:1 groups were generally 1.3-2-fold higher than 50:1 and 1:0 groups, suggesting that 20:1 might be an optimal primer ratio for ssDNA production. It is also notable that dsDNAs were also amplified even in 1:0 groups, which was thought to be the result of imperfect bindings of primers. Nevertheless, our results proved that ssDNA Lockers can be generated via OAA assembly followed with asymmetric PCR. Single strand DNAs can then be harvested and purified by standard PAGE purification assay.

### miRNA Lockers effectively inhibits miRNA function in mammalian cells

To verify the miRNA-inhibiting function of the miRNA Lockers, we tested our system with miR-214, a tumor-metastasis-related miRNA. Previous reports have shown miR-214 can inhibit RNF8 by directly binding its 3’UTR, while RNF8 was reported to promote EMT in cancer cells **(Figure 3a)**. Hence, the inhibitory effects on miR-214 can be observed in multiple ways, including the abundance of miR-214 and its target gene, as well as RNF8 and EMT-related phenotypes such as cell proliferation and migration. Here, we designed two types of miR-214 Locker as shown in Figure 2a. Locker-1(Lc-1) was designed containing two miR-214 binding sites, while Locker-2 (Lc-2) contained 3 binding sites. The control locker (Lc-C) was constructed by replacing the miRNA binding sites of Lc-1 with equal length of poly-C sequence to avoid unnecessary cross talks with other host miRNAs while maintaining the same structure.

**Figure 3.**
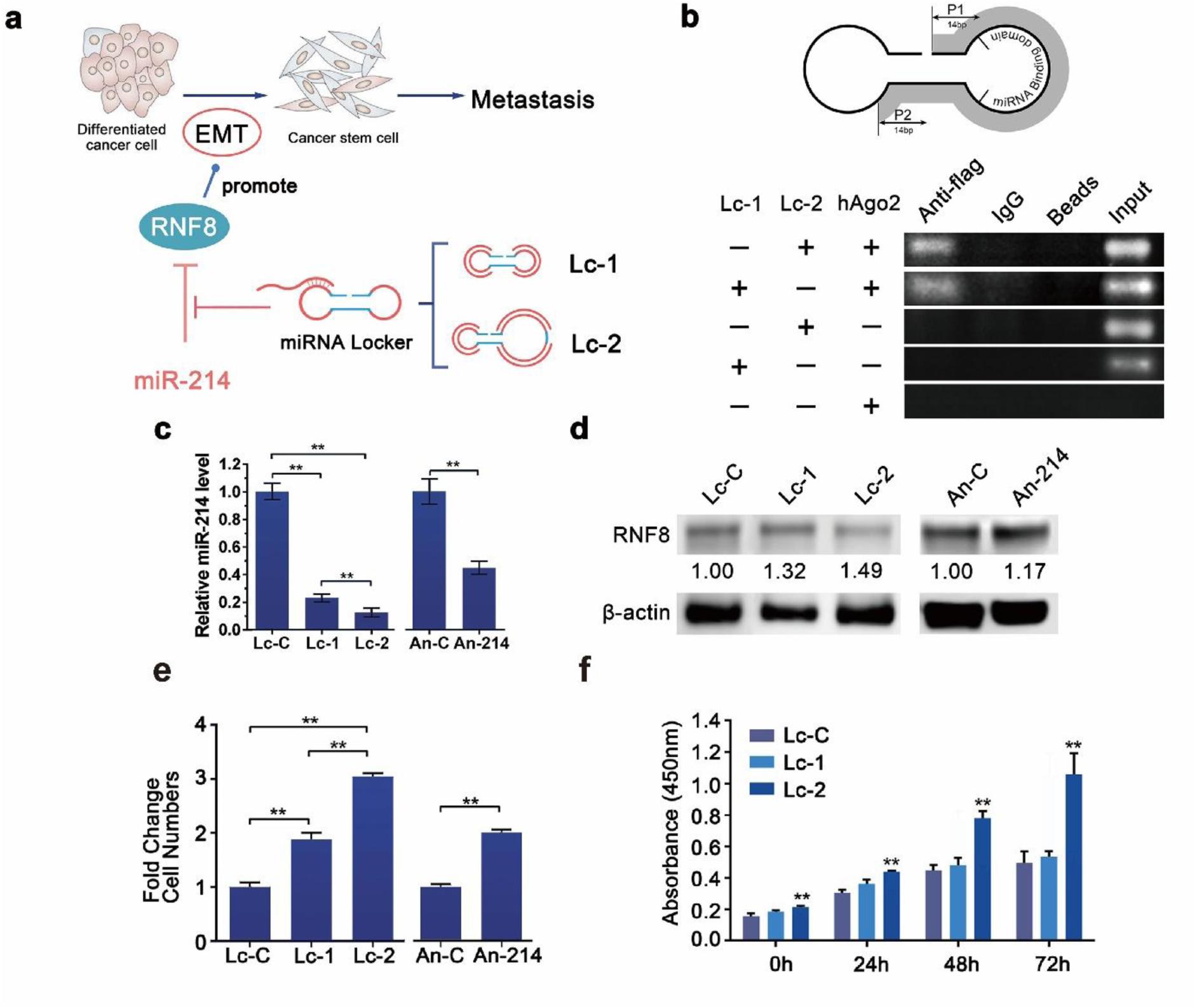
Validation of the miRNA inhibitory effect of miRNA Lockers. (a) Schematic representation showing the effect of miR-214 on EMT process of cancer cells. Two miRNA Lockers, Lc-1 and Lc-2 are expected to promote EMT by blocking the repressive effect of miR-214 on RNF8, A EMT promoting protein. (b) IP-PCR assay determining the binding ability of miRNA Lockers with target microRNA in A549 cells. Schematic representation of primer design is shown above, hAgo2 stands for the Flag-tagged human Ago2 expressing plasmid, input indicates an aliquot of total DNA. Antibodies used for immunoprecipitation are indicated above the lanes. (c) RT-qPCR results demonstrating miR-214 abundance in A549 cells transfected with miR-214 Lockers Lc-C (as control)/Lc-1/Lc-2 or miR-214 Antagomir (An-214)/Antagomir Control (An-C). (d)Western blot determining RNF8 expression level in A549 cells transfected with miR-214 Lockers Lc-C/Lc-1/Lc-2 or Antagomir An-214/An-C. (e) Transwell assay determining of migration in A549 cells transfected with miR-214 Lockers Lc-C/Lc-1/Lc-2 or Antagomir An-214/An-C. (f) CCK8 Assay indicating the proliferation of A549 cells transfected with miR-214 Lockers Lc-C/Lc-1/Lc-2. Relative gene expression was calculated using the 2^-ΔΔCT^ method, with initial normalization of genes against U6 snRNA within each group. The expression levels of each gene in the control groups (Lc-C or An-C) were arbitrarily set to 1.0. Relative protein expression levels were calculated by using β-actin expression level as initial normalization and then set the protein expression level in the control groups (Lc-C or An-C) arbitrarily as 1.0. The relative expression values were averaged from the data in three parallel reactions, and the results were obtained from at least three independent experiments. Error bars represent SD.

To evaluate the miRNA binding ability of the Lockers in living cells, an immunoprecipitation-PCR assay (IP-PCR) was performed to examine the RISC-mediated interaction between the Lockers and its target miRNA. Primers were designed on the supporting module amplifying one of the loop structures of the Locker to minimize the negative effect of highly-repeated miRNA binding sequences on the efficiency of the PCR reaction **(Figure 3b, upper scheme)**. For such an assay, A549 cells transfected with specific sets of nucleotides were cultured for 36 hours and harvested for IP-PCR assay. As shown in **Figure 3b**, ∼80bp (for Lc-1) or ∼100bp (for Lc-2) DNA fragments were amplified from the precipitates of the groups co-transfecting Flag-tagged hAgo2 (Flag-hAgo2) expressing plasmid and Lc-1/Lc-2 by using anti-Flag antibody. By contrast, the negative control immunoprecipitations that did not use antibodies (Labeled as Beads) or adopted the normal rabbit IgG showed no amplification signal. Meanwhile, the control groups transfecting either miR-214 Lockers or Flag-hAgo2 expressing plasmid did not display any amplification signal. These results suggested that miRNA lockers were efficiently recognized by ago-miRNA complexes in *ex vivo* cell cultures.

Then, to evaluate the effect of miR-214 Locker on miR-214 abundance in A549 cells, Taq-man based Real-Time Quantitative Reverse Transcription PCR (RT-qPCR) assay was performed. For such an assay, A549 cells transfected with different Lockers or commercially purchased microRNA inhibitors were cultured for 36 hours before harvesting. Locker Lc-C was used as the negative control for Lockers, while commercially purchased antagomir for miR214 (An-214) was used as the positive control. The Control antagomir molecule (An-C) was used as the control group for An-214. Significant reduction of miR-214 abundance was observed in both Lc-1 and Lc-2 groups compared to the Lc-C group with ∼80%-90% reduction rates **(Figure 3c)**. A similar phenomenon was also observed in antagomir groups with a lower reduction rate (around 60%, **Figure 3c**), which was in accordance with the previous report. Such results suggested a better inhibitory effect of miR-214 Lockers compared to commercially purchased products.

Subsequently, Western blot analysis was performed to examine the effect of miR214 Lockers on the expression level of RNF8, a previously reported gene regulated by miR-214. Results showed that RNF8 levels in the Lc-1 and Lc-2 groups increased by 1.32 and 1.49 folds compared to the Lc-C group, while the RNF8 expression level in the An-214 group increased by 1.17 folds compared to the control group **(Figure 3d)**. Such results demonstrated the capability of miR214 Lockers to affect the expression level of RNF8, a miR-214-regulated gene, in a similar but more effective manner compared to miRNA antagomirs.

Since miRNA inhibitors, such as Antagomirs, were normally utilized for miRNA loss of function research. For further verification, we examined whether miRNA Locker generates miRNA knockdown-correlated phenotype changes identical to those generated by commercially purchased miRNA inhibitors.

Knock-down of miR-214 expression level has been demonstrated to promote the growth of lung adenocarcinoma cells and be involved in migration and invasion of lung cancer via downregulation of RNF8. We then tested if the miR-214 Lockers can induce phenotype changes as expected. For such matter, a cell proliferation assay using Cell Counting Kit-8 (CCK8) was performed to evaluate the effect of miR-214 Locker on cell proliferation. As shown in **Figure 3f**, the absorbance of 450nm (indicating the living cell number) in Lc-2 transfected A549 cells was significantly increased compared to the Lc-C groups, indicating that Lc-2 transfection could significantly strengthen cell proliferation capacity **(Figure 3f)**. Furthermore, Transwell analysis was performed to examine the effect of miR-214 Lockers on the migration of A549 cells. Results showed significant increases in cell numbers migrated through the Transwell chamber in Lc-1 and Lc-2 groups comparing the Lc-C group, while miR-214 antagomir (An-214) showed a similar effect. Cell counts demonstrated a ∼2-fold increase in the Lc-1 group and a ∼3-fold increase in the Lc-2 group compared to the Lc-C group, while a ∼2-fold increase is also observed in the An-214 group compared to the control group **(Figure 3e)**. These results suggested the capability of miR-214 Locker to induce phenotype changes in a miR-214-antagomir-similar pattern.

### miRNA Locker helps determine the effect of miR-654 in the EMT regulation of human lung adenocarcinoma cells

Then, we went on to demonstrate how miRNA Lockers can be used in miRNA research as a promising substitute for current miRNA inhibitors by identifying a new RNF8-targeting miRNA and establishing its regulatory relationship with EMT.

By setting RNF8 as the target downstream gene in three different computational algorithms that analyze miRNA binding sites on the 3’-UTR of a specific gene (TargetScan, miRBase and miRWalk), miR-654-5p (miR-654 for short) was predicted as a promising candidate regulator of RNF8 by directly bind with its 3’-UTR **(Data not shown)**. However, such regulatory relationship has not been experimentally validated yet, while the relationship between miR-654 and cancer metastasis also remains poorly discussed. Hence, we designed four Lockers for miR-654, named as Lc-1p, Lc-1np, Lc-2p and Lc-2np. Locker-1p (Lc-1p) and Locker-1np (Lc-1np) contained two miR-654 binding sites, while Locker-2p (Lc-2p) and Lc-2np (Lc-2np) contained three miR-654 binding sites. Suffix ‘p’ and ‘np’ was marked to distinguish the non-perfect complementary base pairing (np) and perfect complementary base pairing (p) between miRNA Locker and target miRNA **(Figure 4a)**. The control locker (Lc-C) was constructed similarly as the control locker of miR-214 by replacing the miRNA binding sites of Lc-1p with equal length of poly-C sequence to avoid unnecessary cross talks with other host miRNAs while maintain the same structure.

**Figure 4.**
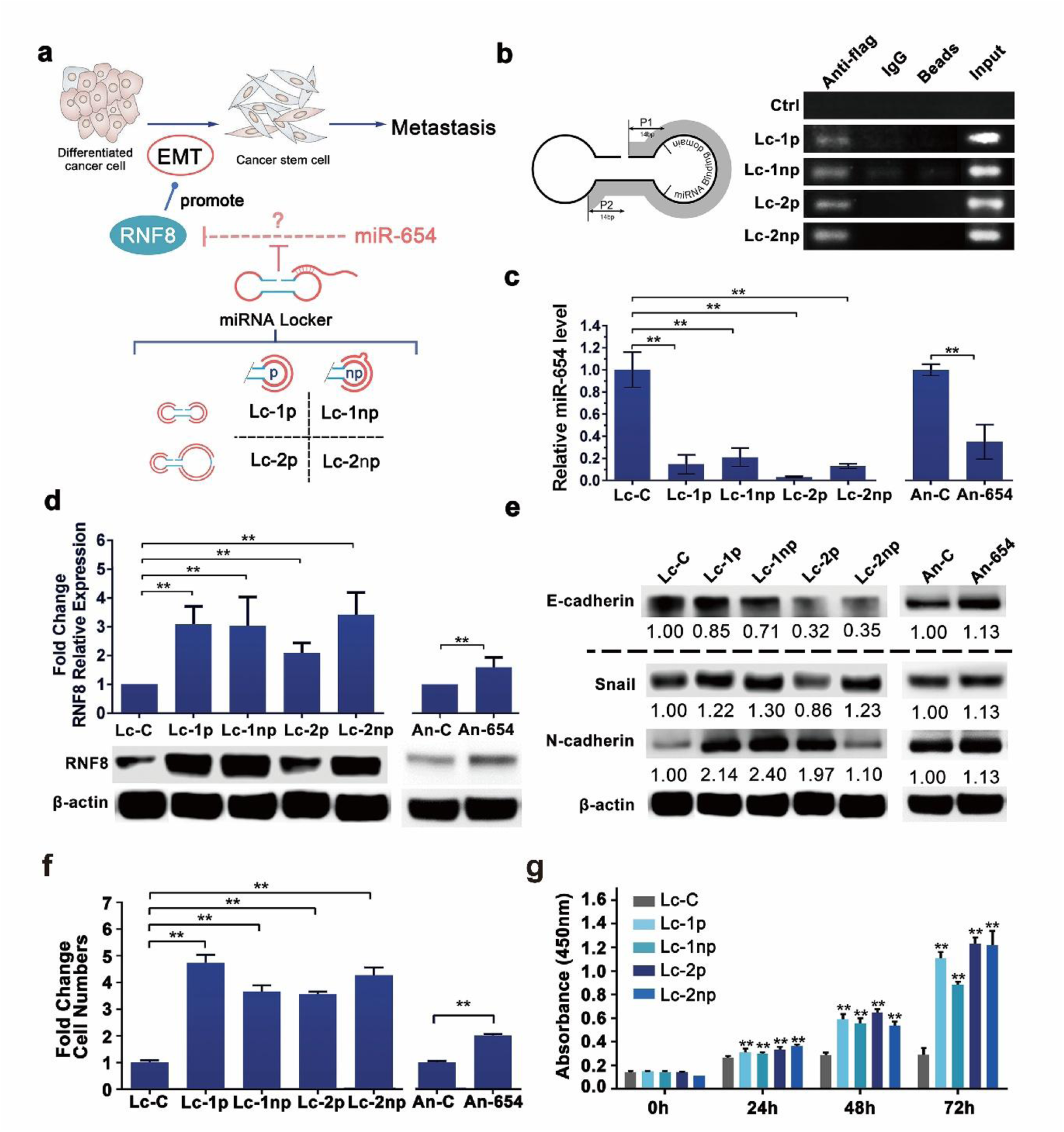
Utilizing miRNA Locker to determine the effect of miR-654 in the EMT regulation of human lung adenocarcinoma cells. (a) Schematic representation showing the anticipated effect of miR-654 on EMT process of human lung adenocarcinoma cells, the relation between miR-654 and RNF8 is unverified. Four miRNA Lockers, Lc-1p, Lc-1np, Lc-2p, Lc-2np are expected to promote EMT by blocking the possible repressive effect of miR-654 on RNF8. (b) IP-PCR assay determining the binding ability of miRNA Lockers with miRNA in A549 cells. Schematic representation of primer design is shown above. Each group was co-transfected corresponding Lockers/Lc-C with hAgo2 expression plasmid. Input indicates an aliquot of total DNA. Antibodies used for immunoprecipitation are indicated above the lanes. (c) RT-qPCR results demonstrating miR-654 abundance in A549 cells transfected with miR-654 Lockers Lc-C/Lc-1p/Lc-1np/Lc-2p/Lc-2np or Antagomir An-654/An-C. (d) RT-qPCR results showing RNF8 mRNA expression level in A549 cells transfected with miR-654 Lockers Lc-C/Lc-1p/Lc-1np/Lc-2p/Lc-2np or Antagomir An-654/An-C. (e) Western blot analysis determining the expression level of epithelial marker E-cadherin and mesenchymal markers (N-cadherin and Snail) in A549 cells transfected with miR-654 Lockers Lc-C/Lc-1p/Lc-1np/Lc-2p/Lc-2np or Antagomir An-654/An-C. (f) Transwell assay determining of migration in A549 cells transfected with miR-654 Lockers Lc-C/Lc-1p/Lc-1np/Lc-2p/Lc-2np or Antagomir An-654/An-C. (g) CCK8 Assay indicating the proliferation of A549 cells transfected with Lockers Lc-C/Lc-1p/Lc-1np/Lc-2p/Lc-2np Relative gene expression was calculated using the 2^-ΔΔCT^ method, with initial normalization of genes against U6 snRNA within each group. The expression levels of each gene in the control groups (Lc-C or An-C) were arbitrarily set to 1.0. Relative protein expression levels were calculated by using β-actin expression level as initial normalization and then set the protein expression level in the control groups (Lc-C or An-C) arbitrarily as 1.0. The relative expression values were averaged from the data in three parallel reactions, and the results were obtained from at least three independent experiments. Error bars represent SD.

As such, IP-PCR was performed to verify the miR-654 binding ability of the designed Lockers. Similarly, primers were designed on the supporting module amplifying one of the loop structure of the Lockers. A549 cells transfected with specific sets of nucleotides were cultured for 36 hours and harvested for IP-PCR assay. As shown in Figure 3b, ∼80bp (for Lc-1p and Lc-1np) or ∼100bp (for Lc-2p and Lc-2np) DNA fragments was amplified from the precipitates of the groups co-transfected with Flag-hAgo2 expressing plasmid and Lc-1p/Lc-1np/Lc-2p/Lc-2np miRNA Lockers by using anti-Flag antibody, while the negative control that did not use antibodies or adopted the normal rabbit IgG showed no amplification signal. Meanwhile, the control group transfected with Flag-hAgo2 expressing plasmid alone did not display any amplification signal. Besides, the knockdown effect of miR-654 Lockers on miR-654 in A549 cells were validated through Taqman-based RT-qPCR assay. Results showed that miR-654 level significantly reduced in four miR-654 Lockers’ groups comparing to the Lc-C group with ∼80%-98% reduction rates **(Figure 4c)**, indicating a good inhibitory effect of miR-654 Lockers on miR-654. Similar phenomenon could be observed on antagomir groups with a lower reduction rate (around 60%, **Figure 4d**). Our results suggested that our fine designed miRNA lockers were efficiently recognized by ago-miRNA complex and further functioned by lowering the target miRNA level.

Since prediction results suggested that miR-654 might downregulate RNF8 expression by directly targeting 3’UTR of RNF8 mRNA. To validate such prediction, Western blot analysis was performed to examine the protein expression level of RNF8 under miR-654 knock-down condition. As showed in Figure 3d, western Blot results showed 2-fold to 3.5-fold increases on RNF8 protein level in Locker-supplemented groups comparing to the Lc-C group, while the RNF8 expression level in An-654 group only increased for 1.17 folds comparing to the control group. In addition, miR-654 knockdown reduced epithelial marker (E-cadhein) and increased mesenchymal marker (Snail, N-cadhein) **(Figure 4e)**, which suggested that Locker-mediated miR-654 knockdown might be related with enhanced EMT via upregulation of RNF8. Taking together, these results demonstrated miRNA Locker-induced knockdown of miR-654 could increase expression of RNF8 and plausibly promote EMT.

With previous results, we could conclude that miR-654 participant in the regulation of RNF8 expression in A549 cells. Since it has been proved that RNF8 participated in cell proliferation and migration in non-small lung cancer cell A549. We anticipate miR-654 might be involved in migration and proliferation of lung cancer by regulating RNF8 expression. To further study the role of miR-654 in A549 proliferation and migration, Cell proliferation assay using Cell Counting Kit-8 (CCK8) was performed to detect the migration capacity of A549 cells transfected with four types of miR-654 Lockers or with Lc-C control. Results showed that the absorbance of 450nm (indicating the living cell number) in four Lockers-groups were significantly increased comparing to the Lc-C groups **(Figure 4g)**, indicating that the knockdown of miR-654 could significantly strengthen cell proliferation capacity. Subsequently, Transwell assay was carried out to test the migration capacity of Locker-mediated miR-654 knockdown groups. Compared with control, miR-654 knock-down dramatically increased the number of migrated A549 cells with 3.5∼5-fold changes **(Figure 4f)**, suggesting that knockdown of miR-654 accelerates cell migration of non-small cell lung cancer A549 cell. Transwell results of miR-654 knockdown by antagomir (∼2-fold change) confirmed such conclusion. Taken together, we found that block of miR-654 using our miRNA lcoker promoted cell proliferation and migration of lung adenocarcinoma cells.

## Discussion

This study presents a significant advancement in miRNA inhibition technology through the development and application of “miRNA Locker,” a novel platform designed using a modular Overlapped Oligo Assembly (OOA) method. The miRNA Locker system produces dumbbell-shaped single-stranded DNA structures that effectively inhibit miRNAs by binding to Argonaute-miRNA complexes, offering enhanced inhibitory efficiency, cost-effectiveness, scalability, and modularity. These findings not only validate the functionality of miRNA Lockers in *ex vivo* models but also address critical unmet needs in miRNA-based therapies and diagnostics.

miRNA dysregulation is implicated in a wide array of diseases, including cancer, cardiovascular disorders, and neurodegenerative conditions, due to its central role in regulating gene expression, cell differentiation, and apoptosis^16,17^. Current miRNA inhibition strategies, including antisense oligonucleotides, sponges, and CRISPR-based depletion tools, often face limitations such as high production costs, off-target effects, and limited scalability^18–20^. The miRNA Locker platform directly addresses these challenges by providing an innovative modular system that combines high specificity and efficiency with cost-effective and customizable production, making it an attractive alternative to existing methods.

Experimental results demonstrated the superior inhibitory efficiency of miRNA Lockers compared to commercially available antagomirs. For instance, miRNA Lockers achieved over 90% reduction in miRNA levels in *ex vivo* lung adenocarcinoma cells, far exceeding the ∼60% inhibition observed with antagomirs. This enhanced efficacy is likely attributable to the high stability of the dumbbell-shaped structure and the precise design of the Lockers, which ensures optimal base-pairing with target miRNAs. These results suggest that miRNA Lockers not only minimize off-target effects but also offer a more robust approach to miRNA inhibition, addressing a critical limitation of current miRNA-based therapies.

Beyond miRNA inhibition, our study demonstrates the ability of miRNA Lockers to induce biologically relevant downstream effects. For miR-214, inhibition via Lockers led to increased RNF8 expression—a key regulator of DNA damage response and cellular homeostasis— consistent with the established regulatory relationship between miR-214 and RNF8. Phenotypic assays further revealed enhanced cell proliferation and migration, mirroring but surpassing the effects observed with antagomirs. Importantly, we identified a novel regulatory role for miR-654 in driving epithelial–mesenchymal transition (EMT) through RNF8 upregulation, expanding our understanding of miRNA-mediated regulation in lung cancer metastasis. These findings highlight the potential of miRNA Lockers as a powerful discovery tool for elucidating miRNA-mRNA regulatory networks in complex biological processes.

The OOA method represents a key innovation in this study, enabling the rapid and cost-effective assembly of functional miRNA Lockers. The OOA platform allows for high-throughput assembly and easy customization of Lockers targeting distinct miRNAs by leveraging a library of chemically synthesized oligonucleotides with pre-designed overhang sequences. This approach significantly reduces production costs and time, facilitating large-scale applications in research and clinical settings. Moreover, the modular design of miRNA Lockers enables multiplex inhibition, opening the door to targeting multiple miRNAs involved in complex disease pathways, a critical step toward advancing precision medicine.

While the results of this study are promising, several challenges remain. The performance of miRNA Lockers in vivo requires further evaluation, including their stability in biological fluids, delivery efficiency, and potential immunogenicity. Additionally, the scalability and robustness of the OOA method should be tested in more diverse biological and industrial settings to ensure its applicability across different miRNA targets and disease contexts. Furthermore, while miRNA Lockers demonstrated superior inhibitory effects compared to antagomirs, the underlying mechanisms contributing to their efficacy, such as enhanced stability or binding kinetics, merit further investigation.

In conclusion, the development of miRNA Lockers represents a transformative step in miRNA inhibition technology, addressing key limitations in existing approaches while offering a versatile and low-cost alternative. Beyond functional miRNA studies, miRNA Lockers hold great promise for advancing translational applications, including miRNA-targeted therapeutics, diagnostics, and systems biology. Future research should focus on *in vivo* validation and expanding the scope of applications, providing a foundation for precision medicine and early disease detection. By enabling precise modulation of miRNA activity, miRNA Lockers have the potential to significantly advance our understanding of miRNA biology and open new frontiers in biomedical research and clinical innovation.

## Materials and Methods

### Cell Culture

MDA-MB-231, MCF7, and HEK293T cells were cultured in RPMI 1640 medium (Hyclone) supplemented with 10% fetal bovine serum (GIBCO) and 100 U/mL penicillin-streptomycin (Hyclone). Cells were maintained in a humidified incubator at 37°C with 5% CO₂ and subcultured upon reaching 70–80% confluence. Cells were detached with 0.05% trypsin-EDTA (GIBCO), washed with PBS, resuspended in fresh growth medium, and seeded at the desired density for subsequent experiments.

### CCK8 (Cell Counting Kit-8) Assay

Cells were seeded in 96-well plates at a density of 2,500 cells per well in 100 µL of complete medium and incubated at 37°C with 5% CO₂ for 12, 24, 48, or 72 hours. At the indicated time points, 10 µL of CCK-8 solution (Dojindo) was added to each well, followed by incubation for 2 hours. Absorbance at 450 nm was measured using a microplate reader to assess cell proliferation.

### Lipofectamine2000 Transfection

Cells were seeded in 6-well plates to achieve 70–90% confluency at the time of transfection. Lipofectamine® 2000 reagent (5–12.5 µL; Invitrogen) and DNA (2.5 µg) were diluted separately in 250 µL Opti-MEM® (GIBCO), mixed at a 1:1 ratio, and incubated at room temperature for 20 minutes. The DNA-lipid complex was added to cells in growth medium without antibiotics and incubated for 12 hours. Medium was replaced with fresh complete medium, and cells were analyzed after 24 or 36 hours.

### Transwell Cell Migration Assay

Cells were resuspended in serum-free medium at 1 × 10⁵ cells/mL after trypsinization, and 300 µL of the suspension was added to the upper chamber of Transwell inserts (8 µm pore size). The lower chamber contained 600 µL of complete medium as the chemoattractant. After 24 hours at 37°C, non-migrated cells were removed with a cotton swab, and migrated cells were fixed with 4% paraformaldehyde for 10 minutes, stained with hematoxylin and eosin, and air-dried. Migration was visualized and quantified under a microscope.

### Western Blotting

Cells were lysed in RIPA buffer containing protease and phosphatase inhibitors (Beyotime) on ice for 15 minutes, followed by centrifugation at 13,200 rpm for 15 minutes. Protein concentrations were determined using a BCA Kit (Beyotime). Equal amounts of protein (60– 80 µg) were resolved on 10% SDS-PAGE gels and transferred to PVDF membranes (Millipore). Membranes were blocked with 5% non-fat milk in PBST for 1 hour, incubated with primary antibodies (e.g., RNF8, β-actin) overnight at 4°C, and probed with HRP-conjugated secondary antibodies. Detection was performed using Pierce™ ECL substrate and visualized on a chemiluminescence imaging system.

### Annealing of Oligonucleotides

Complementary oligonucleotides (10 µM each) were combined in a reaction mixture containing 1 µL of Oligonucleotide 1, 1 µL of Oligonucleotide 2, 1 µL of 10× annealing buffer, and 7 µL of ddH₂O, for a final reaction volume of 10 µL. The mixture was incubated at 95°C for 5 minutes in a thermal cycler, followed by gradual cooling to 4°C at a rate of 0.1°C/second to achieve optimal annealing. The annealed product was stored at -20°C or used directly in subsequent experiments.

### Phosphorylation Modification

Annealed double-stranded oligonucleotides (100 nM) were phosphorylated in a reaction containing 2 µL T4 Polynucleotide Kinase (Takara), 2 µL 10× ligase buffer, and 10 µL ddH₂O. The reaction was incubated at 37°C for 30 minutes and used directly in ligation reactions.

### Ligation

Gel-purified DNA fragments (35 ng) and phosphorylated oligonucleotides (2 µL) were ligated using T4 DNA ligase (1 µL) in a 20 µL reaction containing 2 µL 10×ligase buffer. The reaction was incubated at 16°C for 2 hours, followed by heat inactivation at 65°C for 10 minutes. The ligation product was used directly for bacterial transformation.

### Asymmetrical PCR

PCR was performed using plasmid or digested DNA as the template in a reaction containing 0.5 µL forward primer (5 pmol/µL), 0.5 µL reverse primer, 5 µL 2× Taq PCR Master Mix (Takara), and 3 µL ddH₂O. Different forward-to-reverse primer ratios (1:1, 20:1, 50:1, or 1:0) were tested. The cycling conditions were 94°C for 90 seconds, followed by 40 cycles of 94°C for 20 seconds, 40°C for 20 seconds, and 72°C for 20 seconds, with a final extension at 72°C for 10 minutes.

## Acknowledgments

The work was funded by the National Natural Science Foundation of China (No.31870855) and NUDT Research Program to L.Z..

